# The impact of kinship composition on social structure

**DOI:** 10.1101/2024.01.11.575037

**Authors:** André S. Pereira, Melissa A. Pavez-Fox, Jordan D. A. Hart, Josué E. Negron-Del Valle, Daniel Phillips, Catarina Casanova, Delphine De Moor, Lauren J. N. Brent

**Affiliations:** Centre for Research in Animal Behaviour, University of Exeter, Exeter, EX4 4QG, United Kingdom; Research Centre for Anthropology and Health, Department of Life Sciences, University of Coimbra, 3000-456 Coimbra, Portugal; School of Psychology and Neuroscience, University of St Andrews, St Mary’s Quad, South Street, St Andrews, KY16 9JP, UK; ISCSP, University of Lisbon, 1300 – 663 Lisbon, Portugal; Center for Evolution and Medicine, Arizona State University, Arizona, AZ 85281, USA

## Abstract

The relatedness between group members is a potential driver of variation in social structure. Relatedness predicts biases in partner choice and formation of strong relationships among group members. As such, groups that differ in their percentage of non-kin dyads, i.e., in their kinship composition, should therefore differ in the structure of their social networks. Yet the relationship between kinship composition and social structure remains unclear. Here, we used long-term social and pedigree data from a population of rhesus macaques to investigate the relationship between kinship composition and the connectivity, cohesion, potential for transmission and social differentiation of the social networks of adult female macaques. We found no evidence that the social structure of groups composed of greater proportion of unrelated females differ from that of groups with a lower proportion of non-kin. To investigate this unexpected finding, we built agent-based models parameterised with the empirical data to further explore (1) the expected relationship between kinship composition and social structure and (2) why we did not find such a relationship in the empirical data. Agent-based models showed that kinship composition can influence social structure in populations similar to the one studied, but that this effect may only be detectable with a sample size even larger than ours (19 group-years) and with greater variance in kinship composition (proportion of non-kin varied between 0.830-0.922 in the empirical data). The relationship between kinship composition and social structure might be more apparent when comparing data from species that differ strongly in their social organisation, translating into marked differences in kinship composition. This further emphasises the importance of reporting existing and future kinship composition data to deepen our understanding of the evolution of sociality and highlights the potential of agent-based models to better understand empirical results.

## Introduction

Social structure is the emergent property of the interactions and relationships between group members and a key component of social systems (Hinde 1976; Kurvers et al. 2014; Kappeler 2019). Social structure plays an important role in shaping social and ecological processes, such as a group’s resilience to the death of individuals (Williams and Lusseau 2006), vulnerability to the spread of disease (Otterstatter and Thomson 2007) or the prevalence of parasites (VanderWaal et al. 2013), and propensity to produce advantageous cultural traits (Derex and Boyd 2016). An important goal of socio-ecological research has therefore been to understand what factors underpin its variation (Kurvers et al. 2014).

One such factor might be the kinship structure of groups, i.e., how group members are related to each other. Relatedness mediates the type of benefits individuals might gain from interacting with each other and is thus a key driver of variation in social relationships (Hamilton 1964; West et al. 2007). Cooperating with kin can result in direct and indirect fitness benefits, whereas cooperating with non-kin can only provide direct benefits (Hamilton 1964). Kin selection predicts that, all else being equal, individuals should prefer to cooperate with kin (Hamilton 1964). Indeed, there is a large body of evidence that social mammals bias partner-choice and the formation of strong relationships towards their kin (Smith 2014) and living and interacting with kin can be beneficial (Silk 2007; Widdig 2007; Brent et al. 2017). Since social structure emerges from the patterning of social relationships, the kinship structure of groups should impact social structure.

Kinship structure has two described components: mean relatedness and kinship composition. Mean relatedness is the average relatedness across all dyads in a group (Lukas et al. 2005; Briga et al. 2012; Lukas and Clutton-Brock 2018; Dyble and Clutton-Brock 2020). Kinship composition characterises the to what extent groups consist of kin and/or non-kin, thus complementing mean relatedness with explicit information on the dyadic kinship categories that make up groups (Pereira et al. 2023). So far, research into how kinship structure modulates social structure has focused on mean relatedness (Smith 2014), and has shown, for example that groups with higher mean relatedness have denser networks (Smith et al. 2010) and that members of groups with lower mean relatedness are more likely to form coalitions and to reciprocate grooming symmetrically (Lukas and Clutton-Brock 2018). Yet, it is also important to consider the role of kinship composition on social structure. Mean relatedness and kinship composition capture distinct aspects about how dyadic relatedness scales up to the group level, and they might influence social structure in different ways. For example, two groups that have the same mean relatedness might have different social structures depending on the type and number of dyadic kinship categories present. The social structure of a group that is mostly made up of kin dyads might be more cohesive whereas a group that has some very close kin but many non-kin dyads might be more differentiated and modular.

Here, we quantify the impact of kinship composition on social structure in adult female rhesus macaques (*Macaca mulatta*) from the population of Cayo Santiago, Puerto Rico. To get a comprehensive understanding of what aspects of kinship structure are the most important in driving social structure, we also compare the effect of kinship composition on social structure to that of mean relatedness. The females from Cayo Santiago live in variably sized stable social groups that feature one to multiple matrilines (Pfefferle et al. 2016; Widdig, Kessler, et al. 2016) and are therefore likely to vary widely in their kinship composition, making them the ideal study system to ask this question. Furthermore, their social behaviour, group composition and pedigree have been recorded for multiple generations, allowing us to quantify kinship composition in a continuous way, i.e., as the percentage of non-kin dyads that groups feature, and investigate its relationship with the social structure of multiple groups. The female rhesus macaques from Cayo Santiago form strong relationships with non-kin (Cheng et al. in preparation), but show clear biases in partner choice and relationship strength towards maternal and paternal kin (Widdig et al. 2001; Widdig et al. 2002; Berman 2016; Widdig et al. 2016; Siracusa et al. 2023; Cheng et al. in preparation). This suggests that females from groups with a larger proportion of non-relatives should have fewer partners and fewer strong relationships. As such, we predicted that groups with a greater proportion of non-relatives would have social networks that are less connected, less cohesive and have smaller potential for transmission (Table 1). Relationships between kin are likely consistently strong, as kin not only provide indirect fitness benefits but are also often familiar partners, a strong predictor of partner choice in the study population (Widdig et al. 2001; Widdig et al. 2002; Berman 2016). Non-kin relationships, on the other hand, are likely more variable in the study population, as they range widely from non-existent to strong (Cheng et al. in preparation). As such, we predict that groups with a greater proportion of non-relatives would be more socially differentiated, i.e., have more variation in the number and strength of relationships, compared to groups with more relatives. We used eight social network metrics to quantify social structure, five metrics that quantified connectedness, cohesion, potential for transmission and three metrics that quantified social differentiation (Table 1). To assess the relationship between different aspects of kinship structure and social structure, we repeated all analyses using mean relatedness as the predictor.

**Table 1.** Aspects of social structure quantified in relation to kinship composition. The definition of each of the eight network metrics used, as well as the direction and rationale of their predicted relationships to kinship composition. Here, predictions are framed in the direction where groups contain a greater proportion of non-kin dyads (and lower proportion of kin dyads).

| Aspect of social structure | Social network metric used to assess it | Definition | Prediction for relationship with kinship composition (where higher values represent groups with greater proportion of non-kin dyads) |
| --- | --- | --- | --- |
| **Connectivity, cohesion and potential for transmission** | **Density** | The ratio of the number of relationships observed in the network to the total possible number of relationships that the network could have. Denser networks are more connected. | Negative |
|  | **Diameter** | The shortest number of network edges which information/disease would have to travel through to reach the most distantly connected pair of individuals. Networks with a larger diameter are less connected. | Positive |
|  | **Transitivity** | A measure of the probability that two individuals that have a relationship with a common individual having a relationship themselves. Networks that are more transitive are more clustered. | Negative |
|  | **Mean degree** | The mean number of relationships held by individuals in the network. Larger mean degree indicates that individuals have more relationships. | Negative |
|  | **Mean node strength** | The mean value of all relationships formed by individuals in the network. Larger mean strength indicates that individuals interact more with each other. | Negative |
| **Social differentiation** | **Coefficient of variation of (CV) degree** | The ratio of the standard deviation of degree to its mean. Larger values indicate that networks are constituted by individuals with more variable number of relationships. | Positive |
|  | **CV node strength** | The ratio of the standard deviation of node strength to its mean. Larger values indicate that networks are constituted by individuals with more variable strength. | Positive |
|  | **CV edge strength** | The ratio of the standard deviation of the strength of relationships to its mean. Larger values indicate that social relationships are more variable. | Positive |

## Methods

### Study subjects and location

We studied the rhesus macaques from the free-ranging population of Cayo Santiago, Puerto Rico. This population’s origin dates back to 1938, when 409 macaques were introduced to Cayo Santiago from India, and is currently administered by the Caribbean Primate Research Center. The Caribbean Primate Research Center staff collect demographic data 5 days per week, allowing for accurate birth and death dates. Staff also provide animals daily with commercial monkey chow and *ad libitum* water. There is no predation on the island or regular medical intervention, and leading causes of death are injury and illness (Widdig, Kessler, et al. 2016; Pavez-Fox et al. 2022). Groups in this population are formed without human intervention and consist of one or more matrilines of philopatric females and migrant males (Pfefferle et al. 2016; Widdig, Kessler, et al. 2016). The subjects of this study were the adult (≥ 6 yrs) females of six social groups for which we had detailed behavioural data between 2010 and 2017, totalling 19 group-years (group F: 2010-2017; group HH: 2014, 2016; group KK: 2013, 2015, 2017; group R: 2015-2016; group S: 2011; group V: 2015-2017). We used these behavioural data to build 19 social networks, one per group-year.

### Behavioural and relatedness data

Behavioural data were collected during the working hours of the Cayo Santiago field station, 07:30 and 14:00, using 5 min (group HH in 2016 and KK in 2017) or 10 min (all other group-years) focal samples in which all behaviours were recorded continuously (Altmann 1974). Additionally, the identity of the individuals in proximity (i.e., < 2 m) of the focal subject were registered once every 5 minutes, including at the start and end of the focal (Brent et al. 2013). We collected a total of 5,260 hours of focal samples, averaging 5.39±2.43 hours per female per group-year. For the analyses, we used grooming and proximity events, which are considered socio-positive interactions in primates (Cords 1997; Silk, Alberts, et al. 2006; Silk, Altmann, et al. 2006). We measured grooming events by summing the number of grooming events observed between two individuals. We measured proximity events by summing the number of focals in which members of a dyad were observed in proximity of each other in at least one scan sample. For each group-year, we built a social network in which the nodes were the group-members and the edges were the sum of the grooming and proximity events, thus measuring the occurrence of social events between group-members.

We used the multigenerational pedigree maintained by the Caribbean Primate Research Center (Widdig, Kessler, et al. 2016; Widdig et al. 2017) to calculate dyadic relatedness with the kinship2 R package (Sinnwell et al. 2014). This pedigree was built from demographic data and genetic data that have been continuously collected for the entire population since 1956 and 1992, respectively (Widdig, Kessler, et al. 2016; Widdig et al. 2017). Paternity was inferred from genetic data via parentage analysis, and maternity was inferred from demographic data and confirmed by genetic data via parentage analyses (Widdig, Kessler, et al. 2016; Widdig et al. 2017). We defined pairs of individuals with r ≥ 0.125 as kin and those with r < 0.125 as non-kin. This is the threshold for kin-biases in this population of rhesus macaques (Kapsalis and Berman 1996) and in macaques in general (Chapais and Berman 2004). Even though the population is closed, there is no evidence of inbreeding (Widdig, Kessler, et al. 2016; Widdig et al. 2017). We calculated the kinship composition of each group-year as the proportion of adult female non-kin dyads relative to all adult female dyads. We calculated the mean relatedness of each group-year by dividing the sum of all dyadic relatedness values between adult females by the total number of adult female dyads.

### Statistical analysis

Analysing dyadic social data is challenging because estimates of social relationships are inherently non-independent and uncertain (Weiss et al. 2021; Hart et al. 2022; Hart et al. 2023). Recent work has highlighted that permutations, a statistical method that has sometimes been used to analyse dyadic social data, do not account for non-independence better than parametric regression (Weiss et al. 2021; Hart et al. 2022) and may result in poorly estimated effect sizes in the presence of confounds (Franks et al. 2021). Of the proposed alternatives, Bayesian social network methods are particularly suitable because they explicitly quantify uncertainty around the estimates of social relationships (Hart et al. 2023). Here, we used a Bayesian framework to test our predictions by quantifying the uncertainty in the edge weights of the 19 empirical social networks and by incorporating this uncertainty in Bayesian mixed models that we used to test our predictions.

### Bayesian social networks

To quantify uncertainty in the edge weights of our 19 social networks, we used the R package “bisonR” (Hart et al. 2022; Hart et al. 2023) and fit one edge weight model per group year of network data. For each network, the edge weight models generate a posterior distribution of possible networks that vary on the weight of their edges given the sampling effort of each dyad (Hart et al. 2022; Hart et al. 2023). Dyads with larger observation effort are more likely to have narrower distribution of possible edge weights whereas dyads that were observed less are more likely to have a wider distribution of possible edge weights (Hart et al. 2022; Hart et al. 2023). As such, each sample of the posterior represents a possible “real” social network of the group given the empirical data and observation effort. Crucially, the edge weight models allow us to propagate uncertainty in downstream analyses, including the calculation of network metrics and Bayesian mixed models (Hart et al. 2022; Hart et al. 2023).

To fit the edge weight models, we needed to specify the numerator and denominator of the edges of the empirical networks. We specified the numerator to be the sum between grooming events and proximity, and the denominator to be the number of focal animal samples collected on both dyad members. The edge weights thus consisted of an undirected rate of occurrence of social events per focal sample. As such, we fitted the edge weight model with a “count conjugate” model. We used the function “fit_edge” to fit the “count conjugate” model with the default “count conjugate” priors but, to allow the model to fit, we added a weak beta(0.1,1) prior to the edge component of the model. We confirmed that all edge weight models fitted by checking that the rate of occurrence of social events per focal sample predicted by the model matched the data using the function “plot_predictions”. For each edge weight model, we extracted 200 samples of the posterior probability distribution over the empirical edge weights. bisonR’s edge weight models assume a non-zero probability for all potential edges, even when the existence of an edge is exceedingly small, resulting in fully connected networks (Hart et al. 2022; Hart et al. 2023). As such, to calculate network metrics we needed to set an edge weight threshold that allowed us to differentiate the dyads that likely had an edge from those that did not (Hart et al. 2022; Hart et al. 2023). For each sample of the posterior distribution of the empirical networks, we excluded edges weights that had strength below the minimum non-zero edge weight of the empirical network (Pavez-Fox et al., in revision). We then used the R package “igraph” (Csardi and Nepusz 2006) to calculate each network metric (Table 1) for each of the 200 samples of the posterior distribution of all 19 empirical networks.

### Empirical analysis

We built eight Bayesian mixed models to quantify the relationship between the eight network metrics of social structure and kinship composition, and eight more to quantify the relationship between these same metrics and mean relatedness. We incorporated uncertainty in these analyses by aggregating one sample of the posterior per group-year into 200 datasets. This is, we aggregated the first sample of the posterior of each group-year into one dataset, the second sample of the posterior of each group-year into a second dataset and so forth. Each of these 200 datasets had 19 datapoints, one per group-year, and thus represented one possible set of values for the 19 real networks. We used the function “brm_multiple” from the brms R package to run the same Bayesian mixed model on each of these 200 datasets and then combined the results into a single fitted model object (Bürkner 2017; Bürkner 2018).

We set each model with one network metric as the response variable and the kinship composition of the group as the continuous predictor. We included the number of adult females (group size, here after) as a fixed effect to control for the possibility that social structure was influenced by partner availability (Croft et al. 2008; James et al. 2009). For example, individuals might have more partners in larger groups than in smaller groups just because there are more possible partners available. We included the identity of each group as a random effect to control for potential variation that could be explained by group-level characteristics, such as sex-ratio or inter-group dominance. Ideally, we would have controlled for the nested effect of year within group, because the same group might have different characteristics in different years, but doing so over-parameterised the models and prevented them from converging. We set all models with a Gaussian distribution. We repeated these models but set mean relatedness as the predictor instead of kinship composition.

We used the function “get_prior” from brms to set the default priors for all models. To allow the mean relatedness models that had mean node strength and coefficient of variation in node strength as response variable to converge, we further added a broad and weakly informative normal prior with mean 0 and standard deviation (SD) 10 for the population-level effect (class=“b”). To confirm that the addition of this normal prior was appropriate, we checked that the output of these two models were identical when using a normal prior with double and half of the SD of the prior of the original model. We set adapt_delta=0.999 and max_treedepth=15 to reduce the number of divergent transitions and transitions that exceeded the maximum treedepth, both of which may cause a bias in the obtained posterior samples (Bürkner 2017; Bürkner 2018). When the 95% CI did not span zero, we took this as evidence that the estimated effect of the predictor was systematically different from 0. We confirmed if all models converged by checking that all 200 Rhats per “brms_multiple” model were smaller than 1.05 (Bürkner 2017; Bürkner 2018). Furthermore, we checked one of the 200 models that constituted each “brms_multiple” model to visually confirm that its chains mixed and we used the function “pp_check” to run posterior predictive checks, thus confirming that data generated under the fitted model were comparable to the observed data (Gelman and Hill 2007).

### Agent-based model of the relationship between kinship composition and social structure

We predicted that differences in kinship composition would influence social structure. However, our empirical analyses revealed no effect of the kinship composition of groups in any of the network metrics (see Results). To better understand this unexpected finding, we built two agent-based models to assess if and how kinship composition is expected to affect social structure (model one) and to assess why we did not find a relationship in the empirical analyses (model two) (Figure S1) (Siracusa et al. 2023). In model one, we assessed the effect of kinship composition on social structure in an environment where the only difference between groups was their kinship composition, which we allowed to vary from fully related (100% kin dyads, 0% non-kin dyads) to fully unrelated (0% kin dyads, 100% non-kin dyads). This model helped us visualise the expected effect of kinship composition on social structure given the full possible range of variation in kinship composition. We used model two to see if we could recreate the surprising empirical results and, thus, better understand why we did not find a relationship in the empirical analyses. Similarly to model one, the only thing that differed between groups simulated by model two was their kinship composition, but for model two we limited kinship composition to vary within the limits of our empirical data (that is, with the same range of variation in kinship composition we recorded in the rhesus macaques). To further mirror the empirical data in this model, we limited the number of simulations to sets of 19 groups each.

In both models, we aimed to simulate artificial populations that mirrored the general social organisation, kinship structure and social kin-biases of the Cayo Santiago population. As such, for each simulation, we generated a group of 50 adult females, closely matching the mean number of adult females in real groups on Cayo Santiago (see Results; Siracusa et al. 2023). Each simulated group was structured in *n* kin clusters, each with *m* related adult females, mirroring the way that groups from Cayo Santiago are clustered into matrilines (Pfefferle et al. 2016; Widdig, Kessler, et al. 2016; Siracusa et al. 2023). Dyads of individuals that belonged to the same simulated kin cluster were considered kin and dyads that belonged to different kin clusters were considered non-kin. More generally, this kinship structure is likely common to other mix-related societies as the grouping of unrelated individuals is a possible way that groups arise (Socias-Martínez and Kappeler 2019), which might result in groups featuring independent family lines. We simulated the kin biases observed in Cayo Santiago by parameterising the two agent-based models with the empirical mean probability that kin dyads and non-kin dyads had a social relationship and with the empirical mean strength and SD of those relationships (see Results).

Each simulation consisted of the generation and quantification of one network and proceeded as follows. We started by simulating how many kin clusters there were and how many individuals each kin cluster had. We did this differently for the two models. For model one, first we randomly drew the number of kin clusters from a uniform distribution with range 1 to 50. When we drew one kin cluster, all 50 group members belonged to that kin cluster and were related. When we drew 50 kin clusters, all 50 group members were unrelated as each kin cluster was constituted by only one individual. When the number of kin clusters was *n*, where *n* is between 2 and 49, we drew *n* numbers from a uniform distribution with range 1 to 50. Each of these numbers represented the number of individuals belonging to each of the *n* kin clusters. If these kin clusters added to more than 50 group members, we discarded the clusters that brought the number of individuals to over 50. If the kin clusters added to less than 50, we created a new kin cluster with the difference. For example, if we draw four kin clusters, each with 10, 29, 8 and one individual, the simulated group would add up to only 48 individuals. In this situation, we would create a fifth kin cluster with two individuals, bringing group size to 50. For model two, we found what partitions of 50, i.e., what combination of number and size of kin clusters that add up to 50, resulted in a proportion of non-kin dyads between the minimum and maximum empirical proportion of non-kin dyads (see Results). For each simulation, we selected one of these partitions at random, thus determining the number and size of the kin clusters. This method was suitable for model two because it allowed us to set specific minimum and maximum values of proportion of non-kin dyads that mirrored the empirical data and resulted in a wide distribution of kinship composition between these values (see Results). Conversely, using partitions would not have been suitable for model one because most partitions of 50 result in very large proportions of non-kin dyads, whereas the method we used resulted in a more balanced distribution from fully related to fully unrelated.

In the second step of the simulations we constructed social networks by drawing social relationships. This step was the same for the two models. The probability that each dyad had a social relationship depended on whether they were kin or non-kin and was determined by extracting a random value from a binomial distribution with probability equal to the empirical probability that a kin or non-kin dyad, respectively, had a relationship (see Results). If the extracted value was 1, the dyad was determined to have a social relationship, if it was 0, the dyad did not have a relationship. The dyads that had a relationship were given a relationship strength that depended on the kinship of the dyad members and that was extracted from a beta distribution with mean and SD equal to the empirical values (see Results).

After we generated the social networks, we calculated the same eight metrics as for the empirical analysis. To investigate the relationship between kinship composition and social structure, we ran model one for 100 000 simulations, thus generating 100 000 networks in which we allowed the proportion of non-kin dyads to range from 0.00 to 1.00. We then fitted eight linear models, one model for each network metric as a response of kinship composition, from which we extracted the coefficient of regression to describe the relationship between kinship composition and social structure. To see if we could recreate the empirical null results, we ran model two for 200 sets of 19 simulations each. The 19 simulations mirrored the 19 empirical group-years and the 200 sets of simulations mirrored the 200 samples of the posterior that we incorporated in the empirical analysis and allowed us to gauge the consistency of the results. To describe the relationship between each network metric and kinship composition, we fitted eight linear models, one per network metric, for each of the 200 sets of simulations. Each linear model was set with each network metric as the dependent variable and kinship composition as the predictor and we extracted the coefficient of regression to describe their relationship. A scheme of the steps of the two models can be found in Figure S1.

We confirmed that the models were working as intended by running each model 100 000 from which we calculated the proportion of kin and non-kin dyads that had a relationship, as well as the mean and SD of the strength of the simulated relationships. We confirmed that the means of these values corresponded to the empirical values that informed them (Figure S2).

## Results

### Empirical results

Group-years featured an average of 51.737 ± 17.211 adult females (1452.780 ± 837.535 unique dyads), ranging between 19 adult females (171 unique dyads) and 73 adult females (2628 unique dyads). Kinship composition varied between 0.830 and 0.922 proportion of non-kin adult females (Figure S3). Mean relatedness varied between 0.028 and 0.073. On average, 46.7% of kin dyads within a group-year had a social relationship, i.e., were observed grooming and/or in close proximity to each other. The kin dyads that had a relationship had a mean relationship strength of 0.071 ± 0.079 (mean strength across group-years ± mean SD across group-years). On average, 27.1% of non-kin dyads within a group-year had a social relationship, with a mean relationship strength of 0.027 ± 0.019 (mean strength across group-years ± mean SD across group-years). We used these empirical values to parameterise the probability and weight of relationships in the agent-based models.

Contrary to our predictions, we found no evidence that kinship composition influenced social structure in the female macaques from Cayo Santiago. The social networks of groups that were composed of a greater number of non-kin were not less connected or cohesive, did not have less potential for transmission and were not more socially differentiated than the networks of groups that contained a greater proportion of kin (Figure 1, Table 2). Similarly, we found no evidence that mean relatedness influenced any measure of social structure (Table S1). Regarding the effect of group size on social structure, we found that groups with a greater number of females were more socially differentiated (measured as the coefficient of variation in edge strength), both in the kinship composition and the mean relatedness model (Table S2). We also found weak evidence that larger groups were more transitive when controlling for mean relatedness (Table S2). Finally, we found that all network metrics varied considerably across groups (i.e., the random effect of group ID), indicating that a significant part of the variation in social structure was explained by group-level factors (Table S2).

**Figure 1.**
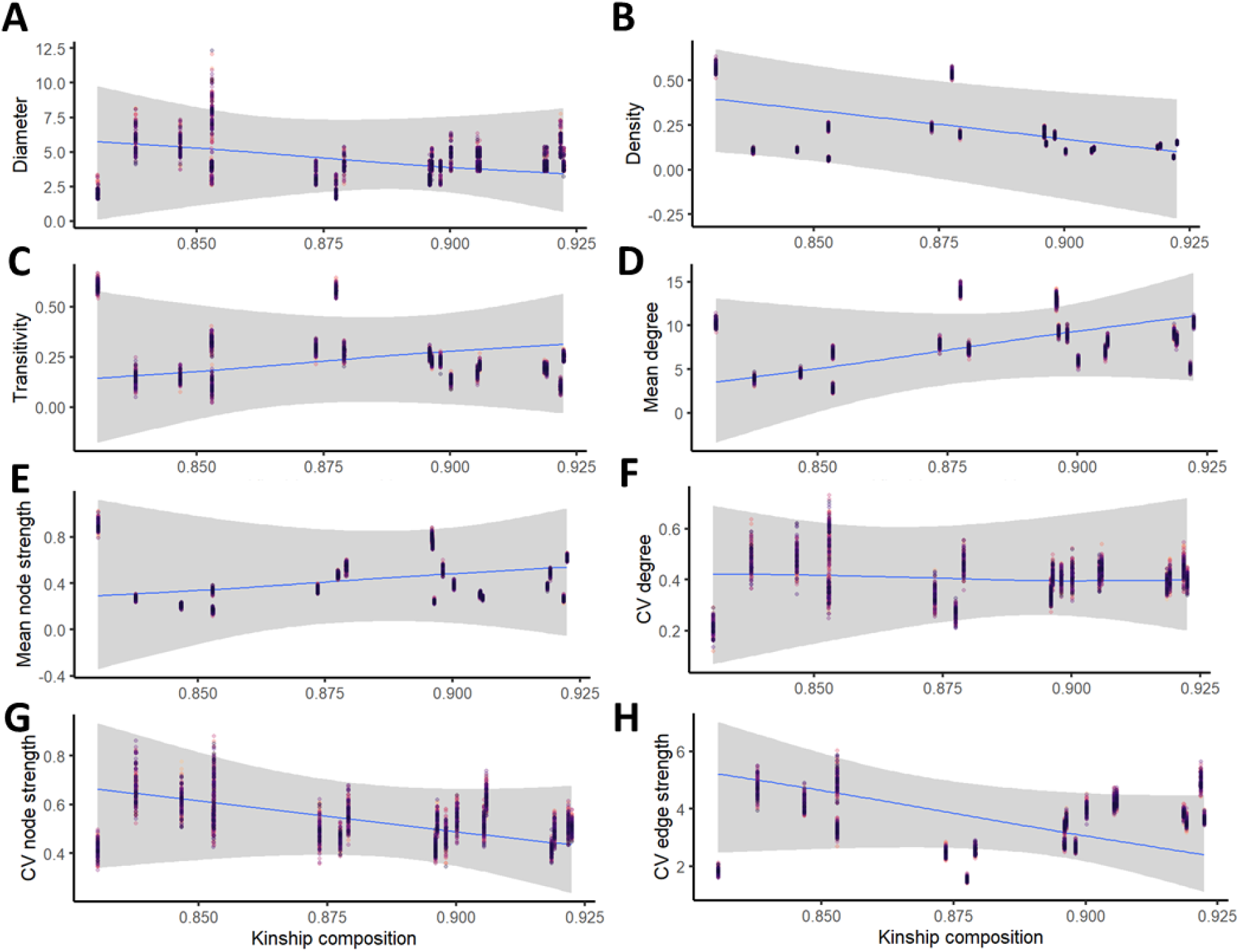
The conditional effect of kinship composition on social structure. The conditional predictions of the regression line between kinship composition and (A) network diameter, (B) network density, (C) network transitivity, (D) mean degree, (E) mean node strength, (F) CV degree, (G) CV node strength and (H) CV edge strength. Group size was fixed to its mean and group ID was accounted in the conditional predictions. Shaded area represents the 95% CI. Points represent measures of each network metric calculated from the 200 datasets that we generated from the samples of the posterior distribution of the edge weight models. As such, in each plot, there are 200 points per group-year. We found no evidence that kinship composition influenced social structure.

**Table 2.** Results from the Bayesian mixed models of the effect of kinship composition on social structure. The 95% CI of the posterior distribution of kinship composition and mean relatedness spanned zero for all network metrics, indicating no effect of kinship composition on social structure.

| Aspect of social structure | Social network metric | Predicted relationship with kinship composition | Result |
| --- | --- | --- | --- |
| Connectivity, cohesion and potential for transmission | Diameter | Negative | Estimate: -20.10; CI: [-89.02; 61.23] |
|  | Density | Negative | Estimate: 1.94; CI: [-3.17; 6.80] |
|  | Transitivity | Positive | Estimate: 1.53; CI: [-4.55; 6.86] |
|  | Mean degree | Negative | Estimate: 75.74; CI: [-71.44; 195.12] |
|  | Mean strength | Positive | Estimate: 2.27; CI: [-9.50; 12.69] |
| Social differentiation | CV degree | Positive | Estimate: 0.07; CI: [-4.39; 4.91] |
|  | CV node strength | Positive | Estimate: -2.35; CI: [-6.73; 2.51] |
|  | CV edge strength | Positive | Estimate: -28.23; CI: [-59.74; 12.20] |

### Agent-based model results

In model one, we set out to characterise the expected effect of kinship composition on social structure given the full possible range of variation in kinship composition. With this model, we generated 100 000 networks that varied from fully related to fully unrelated (Figure S4) and showed that kinship composition influenced some dimensions of social structure to different degrees. In line with our predictions, more unrelated networks were less connected, as they had smaller mean degree and larger diameter (Figure 2). Likewise, these networks were also less dense and less transitive, although the effect of kinship composition on these metrics was not as pronounced (Figure 2). The output of the model showed weak evidence that social differentiation increased with proportion of non-kin dyads and suggested that the relationship between kinship composition and these variables may not be linear (Figure 2). The coefficient of variation in degree and node strength appeared to increase slightly as the proportion of non-kin dyads increased from 0.00 to 0.25, but stabilised afterwards. The coefficient of variation in edge strength appeared to increase as the proportion of non-kin dads increased from 0.00 to 0.50 and to decrease on an accelerating trend afterwards.

**Figure 2.**
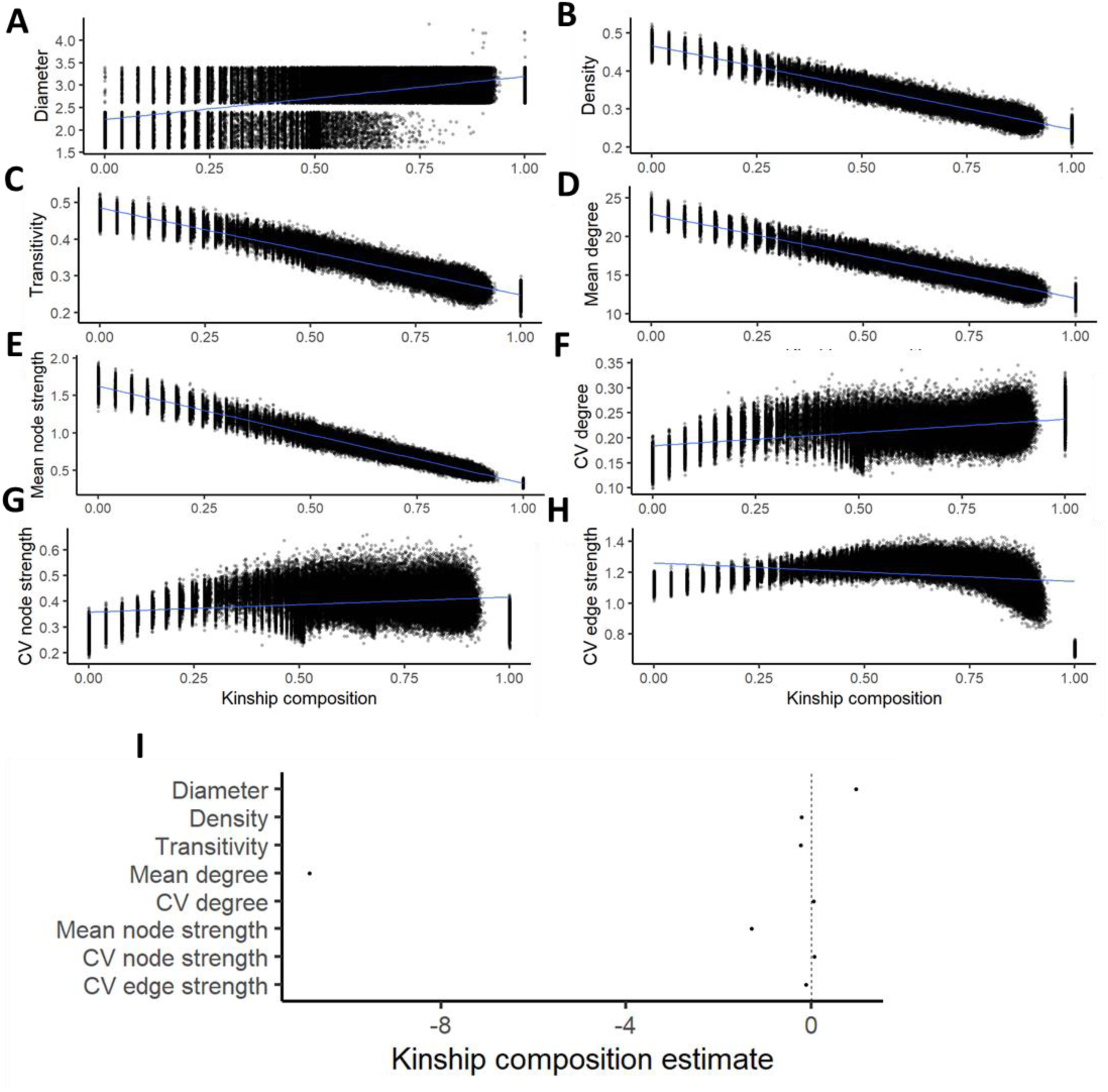
The relationship between kinship composition and social structure in agent-based model one. The relationship between the proportion of non-kin dyads in networks of adult female rhesus macaques and (A) network diameter, (B) network density, (C) network transitivity, (D) mean degree, (E) mean node strength, (F) CV degree, (G) CV node strength and (H) CV edge strength in the 100 000 simulations of model one. Points represent the network metrics calculated in each simulated network. (I) The coefficient of regression of the effect of kinship composition on each network metric from the eight linear models, where values closer to 0 represent a weaker relationship.

We used model two to better understand why we did not find a relationship between kinship composition and social structure in the empirical analyses. In this model, we simulated 200 bouts of 19 networks in which the proportion of non-kin dyads varied from 0.829 to 0.922 (Figure S4). These were the closest values to the empirical variation in the proportion of non-kin dyads that we could generate using partitions of 50. In general, we observed no clear change in social structure as a function of kinship composition (Figure 3). Diameter, density, transitivity, coefficient of variation in degree and node strength showed no clear or pronounced change as the proportion of non-kin dyads increased (Figure 3). Across most simulations, mean degree was smaller in more unrelated networks (Figure 3). However, the relationship between this metric and kinship composition was the most variable, suggesting it might not have been possible to detect a clear signal in the empirical analyses. Mean node strength had a consistently negative relationship with kinship composition and the coefficient of variation in edge strength had a mostly negative relationship, suggesting that we might have been able to detect a relationship between these metrics and kinship composition in the empirical data.

**Figure 3.**
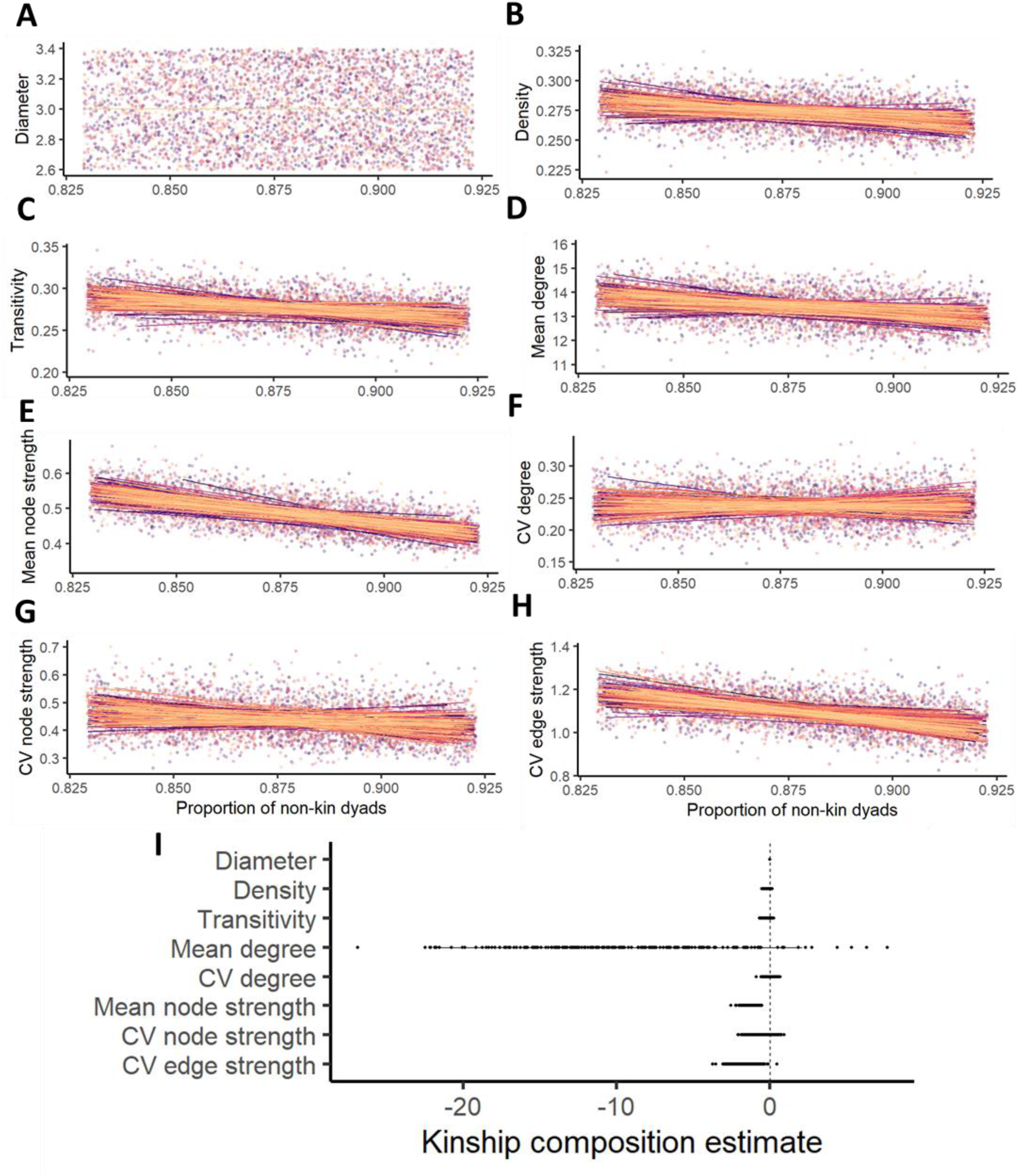
The relationship between kinship composition and social structure in agent-based model two. The relationship between the proportion of non-kin dyads in networks of adult female rhesus macaques and (A) network diameter, (B) network density, (C) network transitivity, (D) mean degree, (E) mean node strength, (F) CV degree, (G) CV node strength and (H) CV edge strength in the 200 “bouts” of 19 simulations of model two. Points represent the network metrics calculated in each simulated network. Data points from the same “bout” were fitted with a linear smoothing term and have the same colour. (I) The coefficient of regression of the effect of kinship composition on each network metric from the eight linear models, where values closer to 0 represent a weaker relationship. Each point represents the regression coefficient of one the 200 linear models that we fitted for each network metric. To visualise the uncertainty in the relationship between each network metric and kinship composition, we plotted the coefficients within the mean ± two standard errors.

## Discussion

We investigated the effect of kinship composition on the social structure of female rhesus macaques. Contrary to our predictions, we detected no evidence of a relationship between kinship composition on social structure. To explore this finding further, we built agent-based models that allowed us to characterise how kinship composition might impact social structure in an environment where only kinship composition was allowed to vary, and to better understand our unexpected empirical result. Overall, the agent-based models mostly confirmed our predictions for the relationship between kinship composition and network structure and helped us better understand our null empirical results.

Kin biases in social behaviour are well known in social mammals (Smith 2014), including in the population of rhesus macaques of this study (Widdig et al. 2001; Widdig et al. 2002; Berman 2016; Widdig et al. 2016; Siracusa et al. 2023; Cheng et al. in preparation). Social structure is the emergent property of the relationships between group members, so we expected that groups with different kinship composition, measured as the ratio of kin to non-kin dyads, should differ in the structure of their networks. In other words, we expected that a network with larger proportion of non-kin dyads, who are less likely to have a relationship and form weaker relationships compared to kin, should, for example, be less dense and have smaller potential for transmission. Yet, we found no evidence that kinship composition influenced the connectivity, cohesion, potential for transmission or social differentiation of networks. Similarly, we did not find an effect of mean relatedness on social structure. Because these results were surprising, we used an agent-based model approach to investigate how kinship composition is expected to affect social structure (model one) and to better understand the results of the empirical analyses (model two) (Siracusa et al. 2023).

Model one showed that, as predicted, kinship composition can influence social structure in a population like Cayo Santiago. The model indicated that groups with a greater proportion of non-kin dyads were less connected, less cohesive and had less potential for transmission than groups with a greater proportion of kin dyads. This suggests, for example, that, in groups that consist mainly of non-kin dyads, information about food sources might take longer to reach everyone; whereas a fully related group might be particularly prone to disease outbreaks because its members are very well connected (Kurvers et al. 2014; Cantor et al. 2021). We did not find a clear relationship between kinship composition and social differentiation. Model one indicated that, as networks went from fully related to featuring some unrelated dyads, degree, node and edge strength became more variable, after which they stabilised, except for edge strength, which decreased afterwards. Overall, model one clearly showed that kin biases at the dyadic level should scale up to affect social structure.

Models one and two indicate that one possible reason why we did not find a relationship between kinship composition and social structure is the limited empirical variation in kinship composition. The models suggest that networks that are very related might have detectable structural differences from networks that are very unrelated, but that such differences might not be detectable between networks that are relatively close in the kinship composition spectrum. This appears to be particularly true for most social differentiation metrics we looked at. The non-linear relationships between kinship composition and network differentiation observed in model one indicate that the ability to detect a relationship in the empirical data might depend on where in the continuum of kinship composition the empirical groups are. It might be easier to detect a difference in social differentiation between a group that is fully related and a group that has 30% non-kin dyads, than between a group that has 30% non-kin dyads and one that has 60%. Even though our data spans 19 group-years, there is not a huge degree of variation in kinship composition in the empirical data and all groups are constituted by a large proportion of non-kin dyads, so they might be less likely to have a real variation in differentiation. The study groups vary in size and number of matrilines, but they share the same overall social organisation, i.e., they are large, female philopatric groups that feature clusters of close kin (Berman 2016; Pfefferle et al. 2016; Widdig, Kessler, et al. 2016), which might limit how much the groups vary in kinship composition. Kinship composition might thus be more likely to differ between groups of species that exhibit markedly different organisations (Pereira et al. 2023). For example, cooperative breeding groups that arise from offspring retention by a single breeding pair might have different kinship composition and kin biases compared to groups that arise from the gathering of independent families. Such contrasts in social organisation might lead to considerable differences in kinship composition and thus to detectable differences in social structure. Furthermore, it is possible that groups with different kinship compositions exhibit differences in kin biases and patterns of association, leading to even larger differences in social structure (Silk 2002; Lukas and Clutton-Brock 2018; De Moor et al. 2020). For example, members of very related groups might not differentiate between related and unrelated partners because the potential of losing indirect benefits is low, resulting in cohesive networks with little differentiation (Clutton-Brock 2006), whereas members of very unrelated groups might exhibit strong kin biases (Silk 2002), resulting in modular and highly differentiated networks. The predicted effect of kinship composition on social structure is evident from model one. As kinship composition and social behaviour data become more widely available in a standardised and comparable way, it will be important to assess how species with different social organisation, behavioural biases and kinship composition differ in their social structure. In the case of the study population, rather than measuring kinship composition as the percentage of non-kin dyads, we might have been more likely to detect differences in social structure if we had measured kinship composition as the number and size of close kin clusters. Different combinations of number and size of close kin clusters can result in the same proportion of non-kin dyads, suggesting that the study population might show more variability in this measure of kinship composition than in proportion of non-kin dyads. We found that group-level effects explained a significant part of the variation across all empirical network metrics. The organisation of groups in clusters of individuals is a key driver of social structure (Kurvers et al. 2014), so the group-level effects might have reflected the way in which each group was organised in close-kin clusters.

The output of model two also suggests that the size of our sample might have limited our capacity to detect an effect of kinship composition on social structure. In general, the slopes of the 200 bouts of 19 simulations varied considerably and were sometimes positive, others negative and others flat. The influence of stochasticity becomes even clearer when we consider that these results are from models in which we have control over confounds and in which the kinship composition of groups is much more evenly distributed than in the empirical world (Figures S3 and S4). The empirical data we used emerged from the collection of individual-level behavioural data from 19 group-years of rhesus macaques and from the maintenance of a long-term pedigree, both involving a notable research effort. Even so, it may still not be enough to detect a relationship between kinship composition and social structure.

The null empirical results might also be explained by variation in group size, which was fixed in the agent-based models but explained a significant amount of empirical variation in some network metrics. Empirical group size and kinship composition were strongly correlated (ρ=0.853; see Supplementary Material), with larger groups being more unrelated (Figure S5). We control for group size in the Bayesian mixed models, which might have further restricted our capacity to quantify the effect that kinship composition has on social structure. Furthermore, the probability that kin and non-kin dyads had a relationship and the mean strength of that relationship decreased as the proportion of non-kin dyads increased (Figure S6), possibly because individuals had to distribute their social availability through more partners (since groups with more non-kin dyads are larger). Overall, these observations further corroborate the relationship between group size and kinship composition (Pereira et al. 2023) and suggest that group size might have explained some important variation in social structure, reducing our capacity to detect a relationship between kinship composition and social structure. Finally, it is also possible that the empirical results corresponded to a true null result. Individuals might be actively adjusting their kin biases to the availability of kin, forming more and stronger relationships with non-kin when they have less kin available (Cheng et al., in preparation). This adjustment could result in groups that have different kin to non-kin ratios having similar social structure.

It is important to note that the empirical and model results we present here only reflect the social patterns of the adult females of the study population. The relationship between kinship composition and the structure of full group networks depends on the kin biases exhibited by adult and juveniles of both sexes and might differ from the relationship we observed and predicted for adult female networks. For example, immature females are known to bias their affiliation towards maternal kin whereas immature, pre-dispersal males are more likely to affiliate with paternal kin (Widdig, Langos, et al. 2016). Mature males can disperse multiple times (Albers and Widdig 2013) and their kin-biases might depend on how many dispersal events they have gone through. The relationship between kinship composition and the structure of full group networks is likely complex and might depend on multiple factors that warrant further investigation. It is also important to recognise that our classification of dyads as “kin” or “non-kin” is likely reductive of the complexity of kin-biases of the study population (Widdig et al. 2001; Widdig et al. 2002; Berman 2016). The relationship between kinship composition and social structure might be more apparent when looking at kinship with a more graded approach.

In summary, we found no evidence that kinship composition influences the structure of the social networks of adult female macaques from the population of Cayo Santiago. Using an agent-based model approach, we illustrated how kinship composition is predicted to affect social structure and found that groups that have a larger proportion of unrelated individuals are less connected, cohesive and have larger potential for transmission of information/disease. Even though the study population seemed to be the ideal system to investigate the relationship between kinship composition and social structure because of the long-term term nature of its pedigree and behavioural data, the agent-based models showed that the sample size and limited variation in kinship composition might have prevented us from finding an empirical relationship. The relationship between kinship composition and social structure might only be apparent when comparing data from species that have markedly different social organisations or when using alternative ways to measure kinship composition. Our study provides a deeper understanding of the interplay between kin-bias, social organisation, and social structure and further highlights the usefulness of agent-based models to complement and better understand empirical analyses (Siracusa et al. 2023).

## Data availability

The data and code necessary to replicate these analyses will be made available once the manuscript is submitted to a peer-reviewed journal.

## Acknowledgements

We are very grateful to Erin Siracusa and the Brent Laboratory for comments and suggestions that increased the quality of this work.

## Supplementary material

### Decision making in building the Bayesian mixed models

Proportion of non-kin dyads (ρ=0.853) and mean relatedness (ρ=-0.732) are correlated with group size. This collinearity can bias the estimates of the coefficients and, while it does not invalidate the analyses, it increases the possibility of finding a false negative. We needed to control for group size (see Methods), so a possible solution would have been to reduce the correlation between the predictor variables and group size. We could have removed the group size factor from the predictor variables. This is, instead of setting proportion of non-kin dyads and mean relatedness as the predictor variables, we could have set number of non-kin dyads and the sum of all dyadic relatedness, respectively.

However, number of non-kin dyads (ρ=0.988) and sum of all dyadic relatedness (ρ=0.975) are even more correlated with group size than the original predictor variables. Furthermore, if we used number of kin dyads and sum of all dyadic relatedness, we would have needed group size to behave as an offset of the response variables, which is not possible to set in Gaussian models. As such, we opted to go with the original, albeit conservative, analyses.

### Supplementary figures

**Figure S1.**
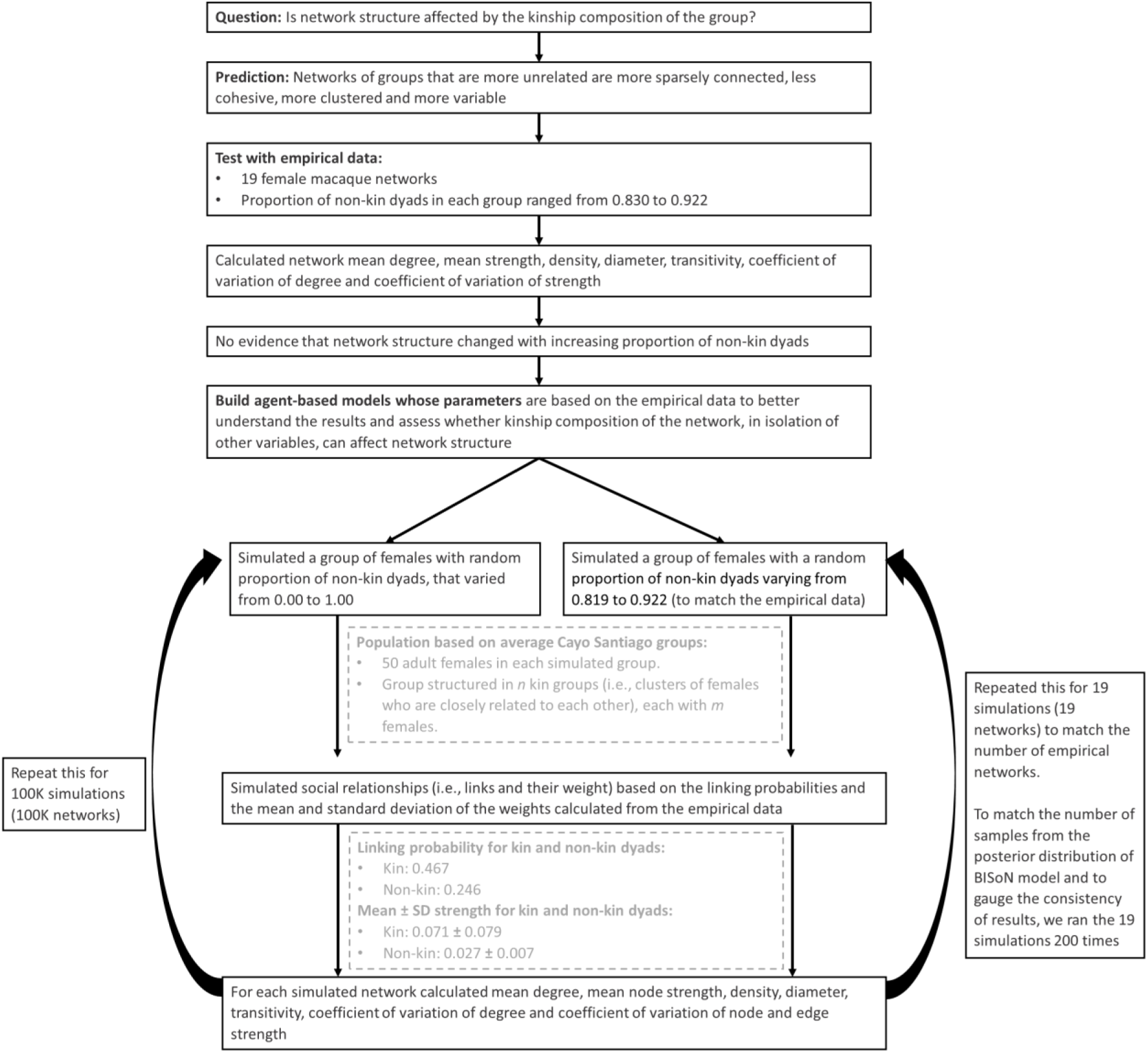
Schematic of the pipeline and of the parameters used in the agent-based model.

**Figure S2.**
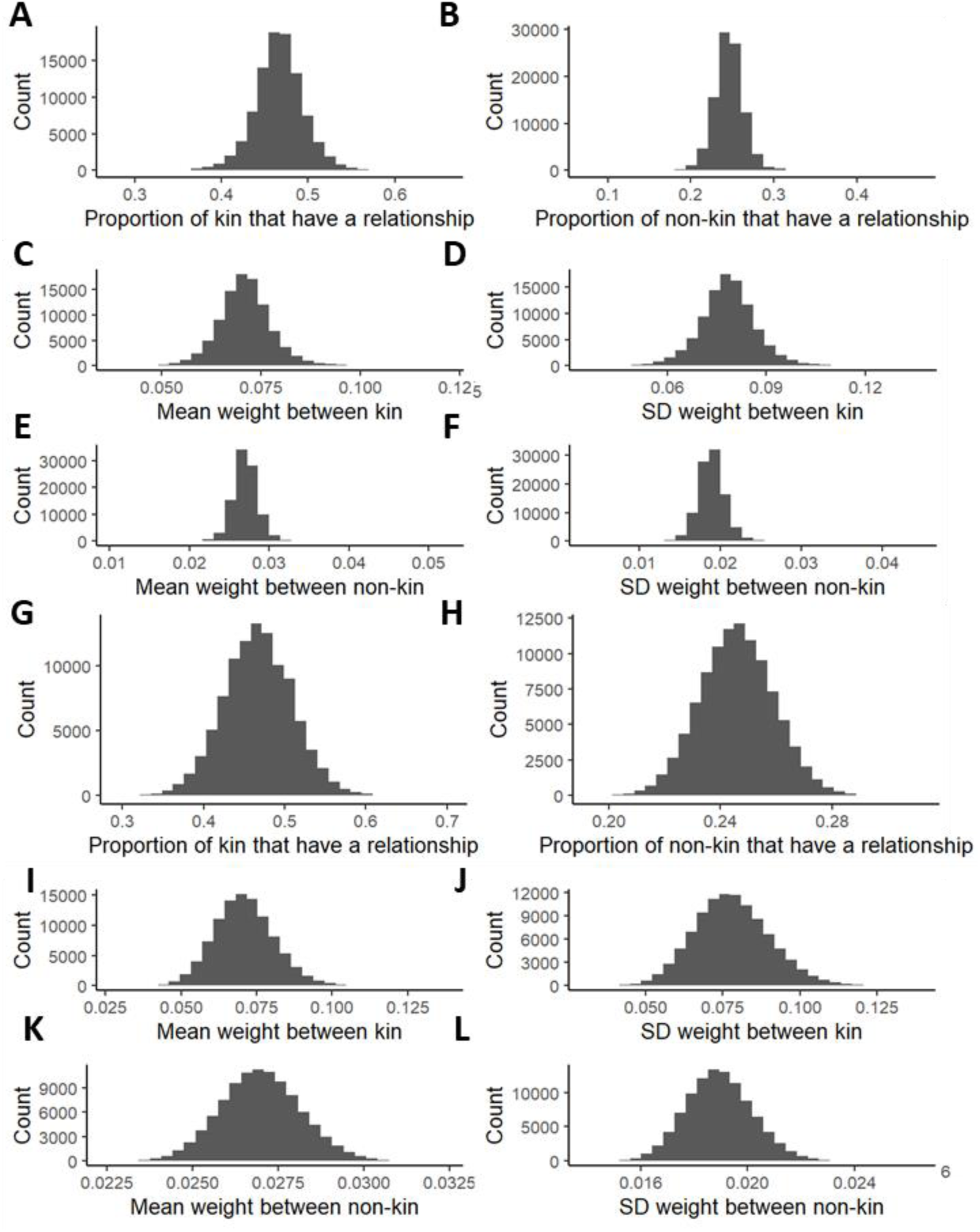
Results from the 100000 simulations from each agent-based models showing the mean proportion of kin and non-kin dyads that had a social relationship and the mean weight and standard deviation of those relationships. For model one, the mean proportion of dyads that had a social relationship was (A) 0.467 for kin dyads and (B) 0.246 for non-kin dyads. The mean ± SD weight of kin relationships was (C) 0.071 ± (D) 0.079 and of non-kin relationships was (E) 0.027 ± (F) 0.019. For model two, the mean proportion of dyads that had a social relationship was (G) 0.467 for kin dyads and (H) 0.246 for non-kin dyads. The mean ± SD weight of kin relationships was (I) 0.071 ± (J) 0.078 and of non-kin relationships was (K) 0.027 ± (L) 0.019.

**Figure S3.**
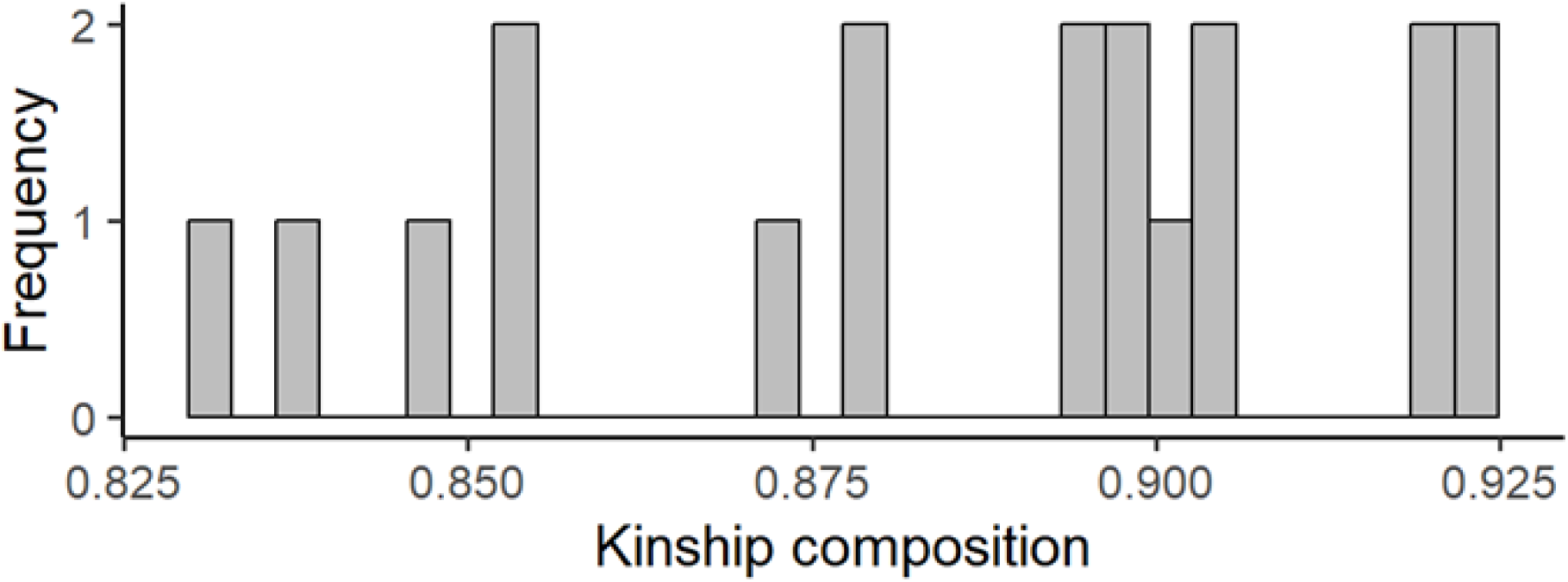
Histogram of the kinship composition (proportion of non-kin dyads) of the empirical group-year.

**Figure S4.**
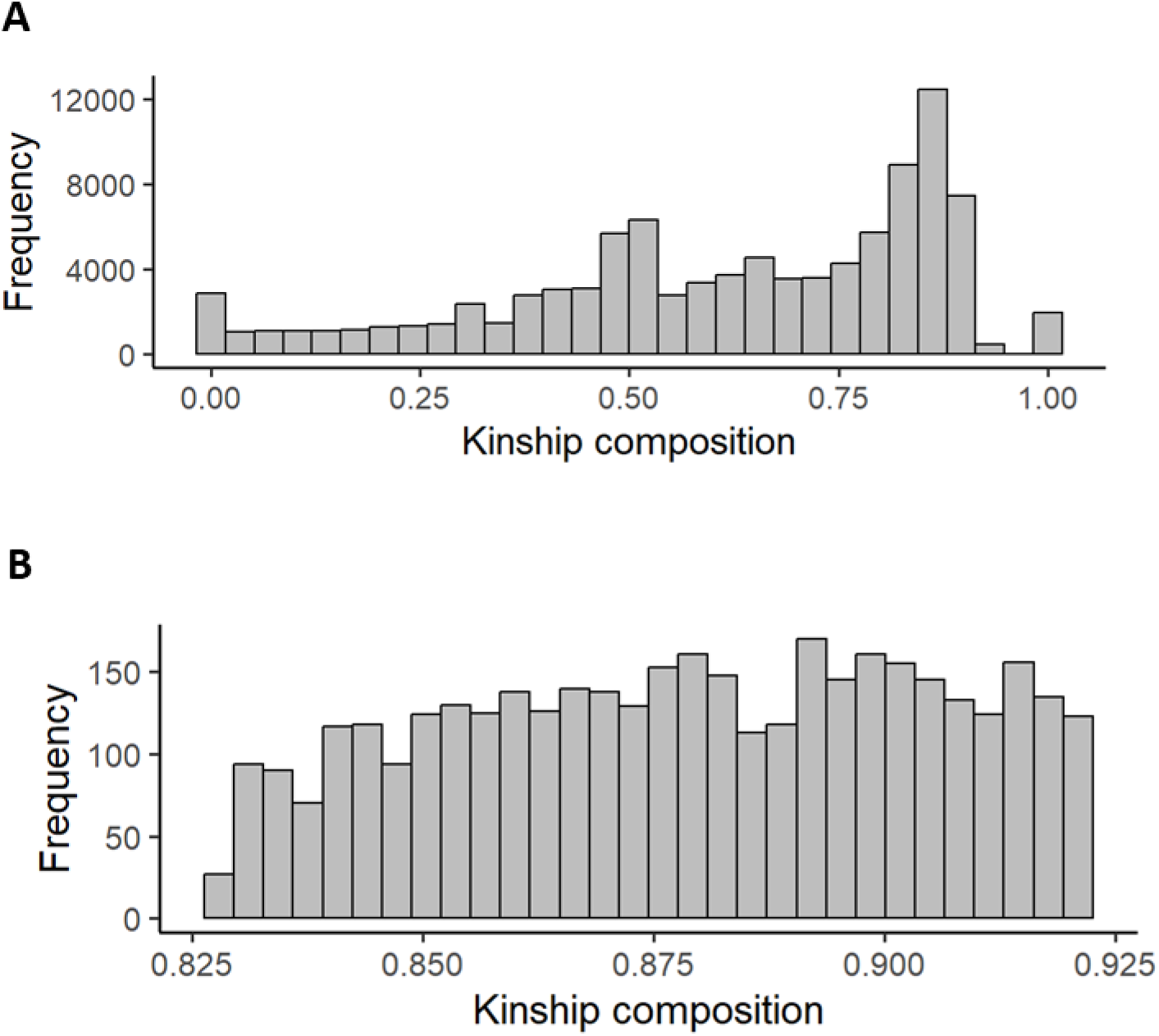
Histogram of the kinship composition (proportion of non-kin dyads) of the networks generated by the agent-based model one (A) and two (B).

**Figure S5.**
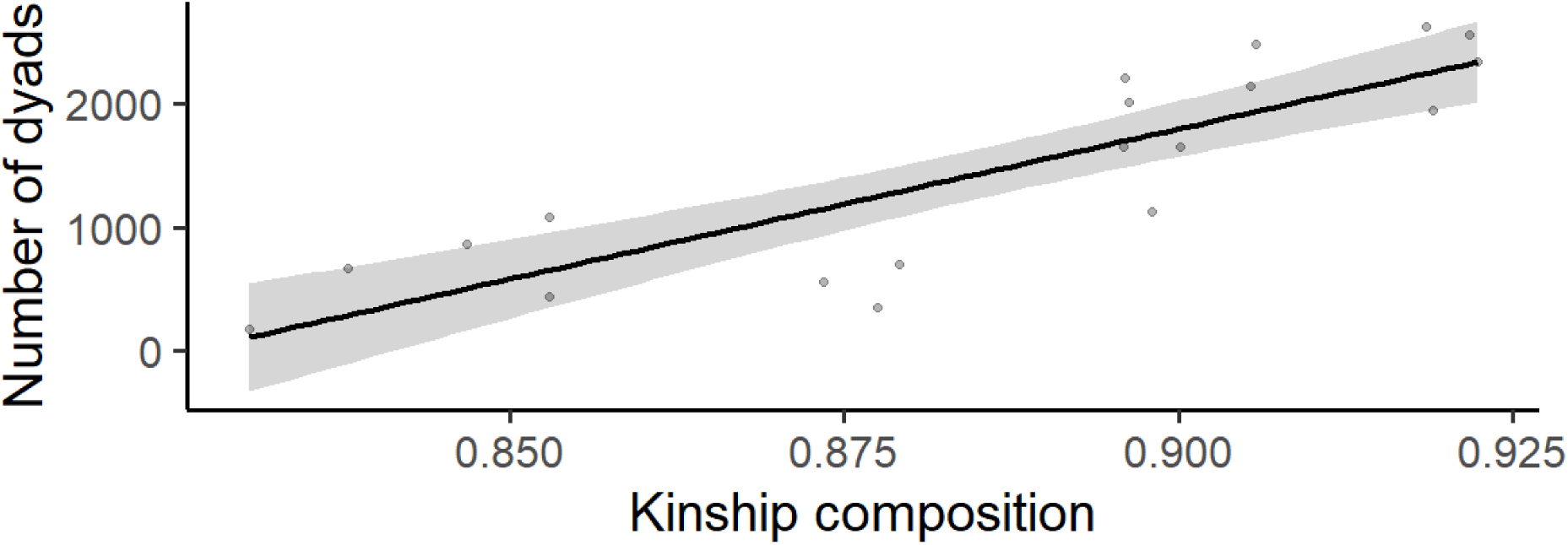
The relationship between kinship composition and number of dyads. The data points represent each of the 19 group-years and were fitted with a linear smoothing term.

**Figure S6.**
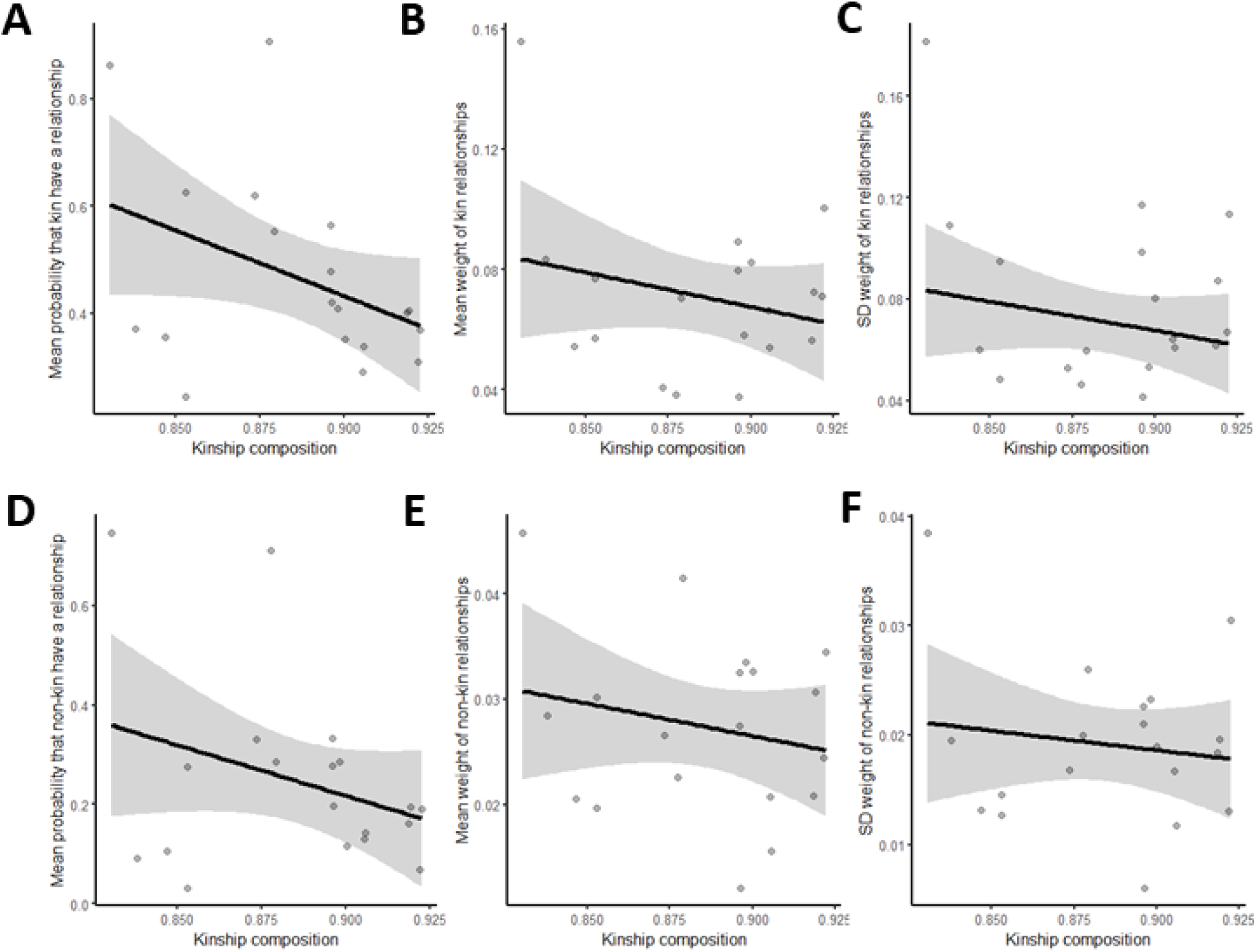
The relationship between kinship composition and the mean probability that kin (A) and non-kin (B) dyads have of having a relationship and the mean (B: kin; E: non-kin) and standard deviation (C: kin; F: non-kin) of these relationships. The data points represent each of the 19 group-years and were fitted with a linear smoothing term.

### Supplementary tables

**Table S1.** Results from the Bayesian mixed models that look at the effect of mean relatedness on social structure. The 95% CI of the posterior distribution of kinship composition and mean relatedness spanned zero for all network metrics, indicating no effect of the mean relatedness on social structure.

| Metric | Prediction | Mean relatedness |
| --- | --- | --- |
| Diameter | Smaller | Estimate: 61.04; CI: [-65.28; 203.32] |
| Density | Smaller | Estimate: -6.48; CI: [-24.18; 6.07] |
| Transitivity | Larger | Estimate: -4.40; CI: [-21.34; 7.72] |
| Mean degree | Smaller | Estimate: -210.78; CI: [-528.26; 43.16] |
| CV degree | Larger | Estimate: 2.04; CI: [-6.65; 11.71] |
| Mean strength | Larger | Estimate: -15.12; CI: [-73.13; 15.82] |
| CV node strength | Larger | Estimate: 1.37; CI: [-5.16; 7.92] |
| CV edge strength | Larger | Estimate: 84.33; CI: [-9.65; 229.82] |

**Table S2.** Estimates and 95% CI for the fixed effect (group size) and for the standard deviation of the random effect (group ID) of the Bayesian mixed models. Terms in bold indicate where the 95% CI did not overlap zero, evidencing that those effects were significant. We took positive 95% CI whose minimum was 0.00, as well as negative 95% CI whose maximum was -0.00 as weak evidence of a relationship, which may or may not be biologically meaningful.

| Metric | Proportion of non-kin dyads |  | Mean relatedness |  |
| --- | --- | --- | --- | --- |
|  | Group size | Group ID | Group size | Group ID |
| Diameter | Estimate: 0.07; CI: [-0.05; 0.18] | **Estimate: 1.33;**<br>**CI: [0.10; 3.43]** | Estimate: 0.07;<br>CI: [-0.01; 0.15] | **Estimate: 1.70;**<br>**CI: [0.37; 3.87]** |
| Density | Estimate: -0.01; CI: [-0.02; 0.00] | **Estimate: 0.14;**<br>**CI: [0.03; 0.37]** | Estimate: -0.01;<br>CI: [-0.01; 0.00] | **Estimate: 0.23;**<br>**CI: [0.06; 0.67]** |
| Transitivity | Estimate: -0.01; CI: [-0.02; 0.00] | **Estimate: 0.14;**<br>**CI: [0.01; 0.38]** | **Estimate: -0.01;**<br>**CI: [-0.01; -0.00]** | **Estimate: 0.20;**<br>**CI: [0.04; 0.58]** |
| Mean degree | Estimate: -0.13; CI: [-0.34; 0.09] | **Estimate: 1.71;**<br>**CI: [0.07; 4.96]** | Estimate: -0.12;<br>CI: [-0.29; 0.04] | **Estimate: 2.74;**<br>**CI: [0.24; 6.80]** |
| CV degree | Estimate: 0.00; CI: [-0.00; 0.01] | **Estimate: 0.13;**<br>**CI: [0.02; 0.35]** | Estimate: 0.00;<br>CI: [-0.00; 0.01] | **Estimate: 0.14;**<br>**CI: [0.04; 0.37]** |
| Mean node strength | Estimate: -0.01; CI: [-0.03; 0.01] | **Estimate: 0.23;**<br>**CI: [0.01; 0.68]** | Estimate: -0.01;<br>CI: [-0.02; 0.01] | **Estimate: 0.47;**<br>**CI: [0.03; 1.72]** |
| CV node strength | Estimate: 0.00; CI: [-0.00; 0.01] | **Estimate: 0.08;**<br>**CI: [0.00; 0.26]** | Estimate: 0.00;<br>CI: [-0.00; 0.01] | **Estimate: 0.11;**<br>**CI: [0.01; 0.32]** |
| CV edge strength | **Estimate: 0.08; CI: [0.02; 0.13]** | **Estimate: 0.58;**<br>**CI: [0.03; 1.77]** | **Estimate: 0.06;**<br>**CI: [0.02; 0.10]** | **Estimate: 1.52;**<br>**CI: [0.36; 4.09]** |

